# Imputing relative abundances via Gaussian random fields

**DOI:** 10.64898/2026.10.12.758512

**Authors:** Andrey Shternshis, Prashant Singh, Luisa W. Hugerth

## Abstract

Longitudinal studies with human participants frequently face the challenge of missing samples, which lead to incomplete trajectories of participant biomarkers and a reduction in data size. Existing approaches focus on single-variable imputation or model features one at a time, while neural networks typically require a large number of parameters and substantial training data. To address these challenges, we present an explainable model for multiple imputation of compositional samples based on a Gaussian random field (GRF). The model is specifically designed for longitudinal data: it incorporates participant-specific information and accounts for both temporal dependence across repeated observations and within-sample correlations, while using a moderate number of parameters. By coupling the GRF with sparse activation functions, the framework outputs sum-constrained compositions and explicitly models zero inflation. Under the Aitchison and Jensen–Shannon distances, our framework achieves imputation accuracy comparable to that of an observation-carrying baseline, while overcoming its static nature by modeling temporal trajectories and multivariate correlations. The resulting multiple imputations allow us to reconstruct complete datasets for downstream analysis and uncertainty quantification. We demonstrate the practical utility of our framework on gut microbiome datasets, restoring incomplete time series and achieving a balanced accuracy of 0.72 in predicting food allergy in infants (sensitivity 0.8) and of 0.64 in forecasting postpartum depression.

## 1 Introduction

Missing samples are a common challenge in longitudinal studies involving human participants. Key causes include missed appointments, insufficient sequencing reads, instrument failure, sample loss during transport, and degradation during storage. Such missingness fragments the longitudinal trajectories of participants’ biomarkers and interferes with the understanding of the underlying temporal dynamics. It also weakens statistical power, making it harder to distinguish true biological trends from random variation. Furthermore, unaddressed missingness risks introducing systematic bias into downstream inferences, as incomplete datasets may fail to represent the overall population. Data loss is especially problematic given that human longitudinal studies are frequently restricted to small or moderate cohort sizes. This limitation is driven by the costs and challenges of recruiting and retaining participants [1], as well as by the attrition that occurs in longitudinal cohorts [2].

Incomplete data are especially common in longitudinal studies, as some participants cannot be measured at every time point: only 9% of reviewed randomized controlled trials with longitudinal measurements reported no missing data [3], while the average retention rate across longitudinal cohort studies was 73% [4]. Sampling in longitudinal microbiome studies may likewise be irregular, with participants having different numbers of observations across follow-up time points [5–7]. Similar issues arise in other areas of biomedical research. A review of molecular epidemiology studies found that 66% had missing data on at least one biomarker or key variable of interest [8]. Similarly, a systematic review of cohort studies with repeated exposure measurements found substantial missing data across the studies examined [9]. Missing data are also an important challenge in multi-omics analysis, where measurements from one or more molecular layers may be unavailable for some samples [10].

One strategy for overcoming data loss is imputation, which predicts missing measurements to recover complete participant trajectories. Rather than discarding incomplete cases and further decreasing the data size, imputation preserves statistical power and temporal continuity across the cohort. However, imputing longitudinal biological data is particularly challenging, because models must capture participant-specific temporal dynamics while generating entire samples that satisfy the strict mathematical constraints of the underlying data type. This work presents a model designed to replace missing *compositional* samples, which represent relative fractions of a whole. Compositional data are encountered across biological omics studies. In sequencing-based transcriptomics, the measured abundance of each gene or transcript is expressed relative to the total amount of sequencing data [11]. Similarly, untargeted proteomics and metabolomics commonly rely on relative quantification of proteins [12] or small molecules rather than on direct measurements of their absolute concentrations [13]. In microbiome studies, sequencing produces a sample-dependent number of reads that reflects the relative representation of the detected taxa. Consequently, the observed data provide information about the relative abundances of taxa, while their absolute abundances cannot be recovered from sequencing data alone [14]. The resulting relative abundances form a composition that describes how the observed microbial community is distributed across taxa, making this compositional structure an inherent property of microbiome data [14, 15].

The central focus of this work is *imputation accuracy* : the aim is to decrease the distance between real samples and samples predicted by the model, thus generating more realistic samples that capture the dynamics of the data. The presented model is stochastic; it therefore allows us to perform *multiple imputation* [16] and to assess the uncertainty in the estimated missing samples [17]. Since the true values of missing data cannot be determined with certainty, generating multiple variants of each sample reflects the natural range of plausible compositions better than relying on a single deterministic guess. In practice, analyses can then be run independently on each of an ensemble of completed datasets, with the final results aggregated so that inferences drawn from the data account for the uncertainty due to missingness.

We propose an interpretable Gaussian random field (GRF) framework. The model is specifically designed for datasets that are typically not large and therefore uses a restricted number of parameters, maintaining an appropriate parameter-to-sample ratio and ensuring that the model can be reliably estimated from the available data. It exploits all available samples during training and operates on a latent space that explicitly captures within-sample covariance structures, temporal autocorrelation, and participant covariates. Because the GRF is inherently probabilistic, it enables multiple imputation, allowing us to propagate imputation uncertainty directly into downstream analyses. The model is interpretable owing to its explicit architecture, which provides greater transparency into the mechanisms underlying its predictions, and it requires neither a dataset-specific architecture nor massive sample sizes.

A common approach for imputing missing values in biological applications is Multivariate Imputation by Chained Equations (MICE) [18]. MICE iteratively imputes each variable by fitting a separate conditional model that uses the remaining variables as predictors. A more recent approach, adversarial random forests [19], combines ideas from generative adversarial networks [20] with ensembles of decision trees [21]. Random-effects models [22] are used to impute normally distributed outcomes in longitudinal studies; while this approach allows for multiple imputation, it still focuses on single-variable imputation. A multivariate normal distribution was proposed for imputation in [23]. A sum of Gaussian processes is used in [24] to model features one at a time. In contrast, modern generative models such as diffusion models [25] and variational autoencoders [26] can model high-dimensional joint distributions, but their neural network architectures typically require a large number of parameters and substantial training data. N-mixture models [27] are designed for count data in ecological studies and can incorporate spatial structure by taking into account the proximity of adjacent positions. Gaussian processes [28] are used to analyze the variance of samples in longitudinal data in [29] and to infer temporal structure in [30]. A GRF coupled with a Dirichlet distribution [31] is used in [32] for inference over high-dimensional categorical observations and in [33] for the imputation of spatial compositional data. To use GRFs in our work, we associate compositional samples with positions defined by participant and time (Fig. 1). When microbiome samples are collected at high temporal frequency, and the time series are therefore long, missing samples can be interpolated by linear regression, nearest-neighbor approaches, differential equations, or Bayesian networks [34], as was done in [7]. However, as longitudinal studies involving human participants often do not include long time series, interpolating from nearby available samples or weighting them is not always possible.

**Figure 1:**
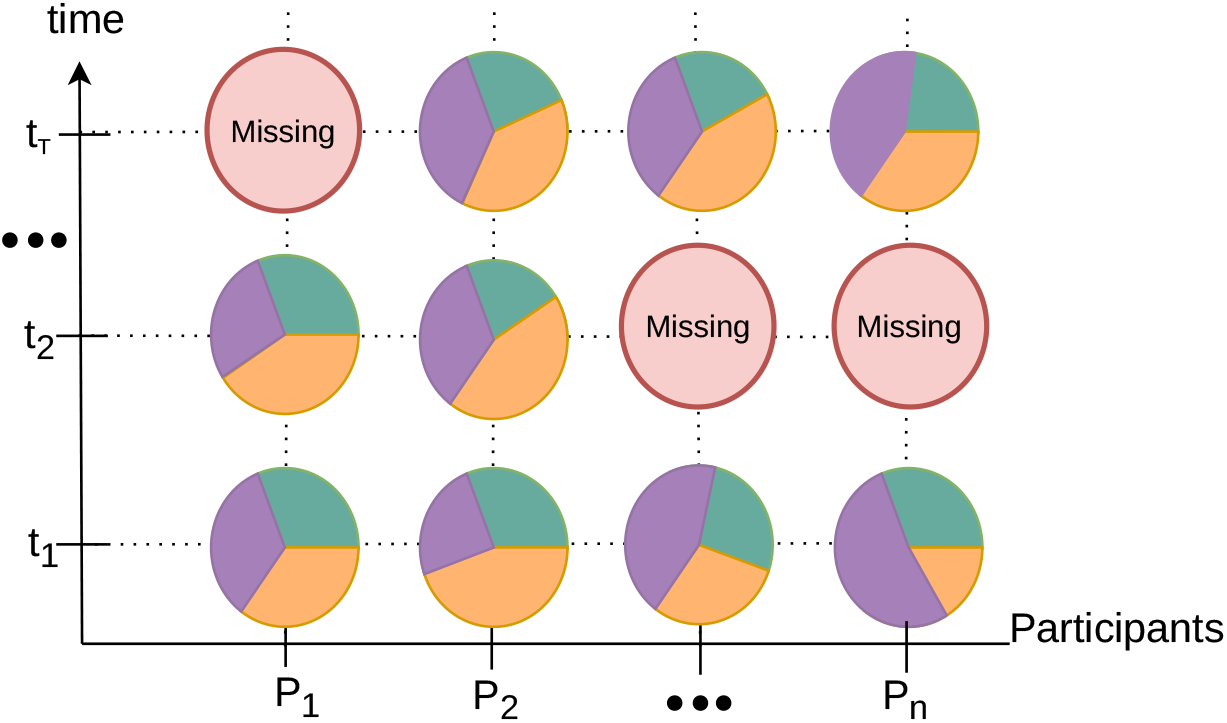
Schematic illustration of a longitudinal study, where *t* denotes time points and *P* denotes participants. Pie charts represent collected samples; some samples are missing.

The rest of the paper is organized as follows. We describe the model used for imputation, the details of its implementation, and the metrics used to evaluate model performance in Section 2. The datasets used for the application are described in Section 3. The main results are presented in Section 4, and Section 5 concludes the paper. Appendix A provides additional experiments on the classification of imputed datasets.

## 2 Methods

This section is organized as follows. We introduce the notation and the arrangement of samples used by the model in Section 2.1. We present the model architecture and describe it in detail in Section 2.2. We give the details of model training in Section 2.3. The metrics for evaluating imputation accuracy are given in Section 2.4, and the pipeline for downstream forecasting, together with its accuracy metric, is described in Section 2.5.

### 2.1 Data representation

All available samples are ordered chronologically for each participant, as shown in Figure 1. The pie charts represent compositional vectors of size *D* (*D* = 3 in the sketch). Each sample has its own position *s* = (*i, j*), that is, a participant *P*_*i*_ and a timestamp *t*_*j*_ . There are *n* participants and *T* time points in total. In addition to the collected samples *y*(*s*) = (*y*_1_(*s*), *y*_2_(*s*), …, *y*_*D*_ (*s*)), participants have additional information *β*, called covariates; the covariate vector *β*(*s*) has the same size *n*_*β*_ at every position. Since some samples are missing, the total number of samples *N* is less than *nT* .

Treating undetected components as present at low abundance avoids zero values, which would otherwise complicate analyses, including data transformations and metric calculations. We therefore represent each component by a positive value, potentially of very low abundance, *y*_*d*_(*s*) *>* 0. This reflects the assumption that an undetected component may still be present but remain below the detection threshold, as the measurement process does not necessarily capture all components present in a sample. We identify the minimum non-zero value across all samples and replace all zero values with half of this minimum, denoted *ϵ*, while ensuring that the sum remains equal to one:

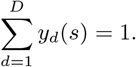

### 2.2 Model for imputation

As a benchmark for comparative evaluation, we include a standard baseline method, “last observation carried forward” (LOCF) [35]. LOCF fills missing entries by carrying over a participant’s most recent sample, or their temporally nearest sample if no prior data exist. While LOCF directly preserves real data values, it serves primarily as a deterministic reference point, as it can neither perform multiple imputation nor account for underlying temporal variability.

A sketch of the proposed GRF model is presented in Figure 2. The core of the model is a latent space in which the variables follow a multivariate normal distribution. The mean values of these variables are defined by the information available for each participant and the timestamp at which the sample was collected. The covariance between variables is based on two components: how features within one sample are correlated, and how the sample relates to other samples from the same participant. The closer the samples are in time, the more correlated they are. Latent variables live in real space, with no restrictions on their values. The model outputs compositional samples by means of activation functions that transform real vectors into compositional vectors. We apply the standard softmax function [36] as well as its generalization, entmax [37], which allows the model to reflect data sparsity, i.e., features equal to zero in compositional space. Compositions that replace missing samples, or that are used to validate the model, are predicted via the posterior distribution known for Gaussian processes [38].

**Figure 2:**
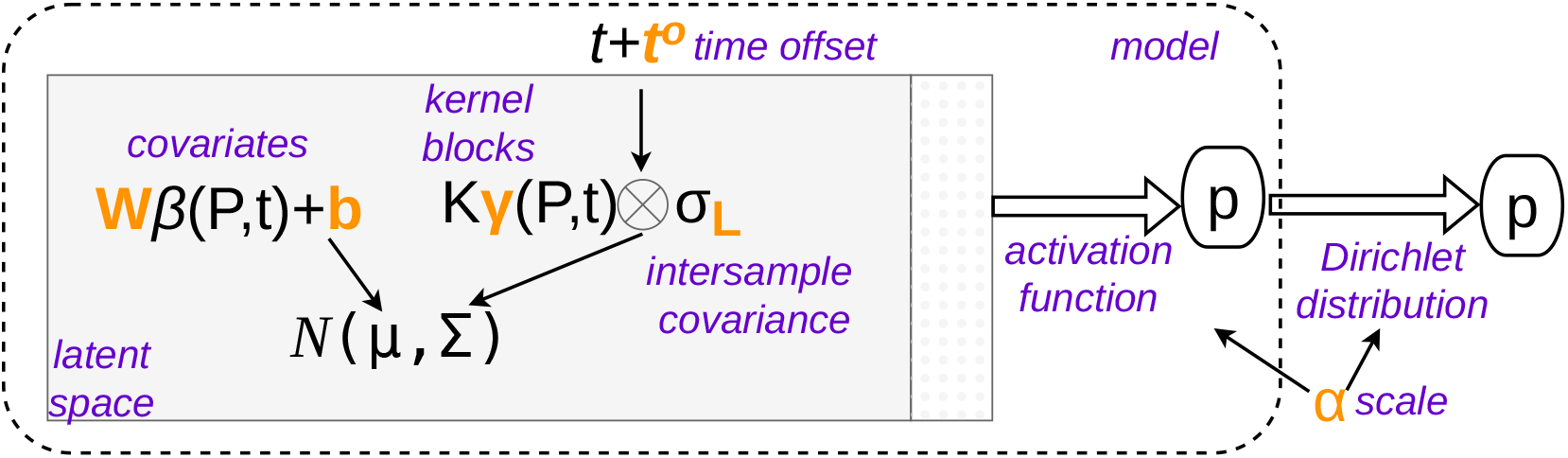
Model for the imputation of compositional data. Inputs to the model are the sample positions (*P, t*). Model parameters are shown in bold (orange). The latent space, shown as a shaded (gray) rectangle, follows a Gaussian distribution, *N*(*µ*, Σ). Compositional data are modeled by applying an activation function to the latent space extended with the last value per sample (dotted column). The output, one compositional sample per position, is depicted as a circled *p*. The optional components, scaling by *α* and sampling from a Dirichlet distribution, are shown outside the dashed region. Brief descriptions of the functions (hollow arrows) and parameters are given in italics (purple).

A complete description follows.

#### Latent space

Each position *s* has a latent vector *z*(*s*) = (*z*_1_(*s*), *z*_2_(*s*), …, *z*_*D−*1_(*s*)) of size *D −* 1. Jointly, all latent vectors follow a multivariate Gaussian distribution:

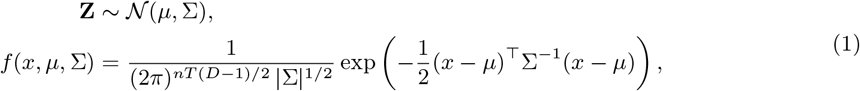

where **Z** = [*z*(1), *z*(2), …, *z*(*nT*)] is the concatenation of all latent vectors and *f*(*x, µ*, Σ) is the density function of the Gaussian distribution. The mean *µ* is defined by a linear transformation of the covariates available for each participant and time point, *β*(*s*):

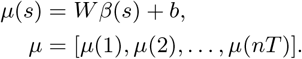

The covariance matrix Σ takes into account both the covariance of features within a sample, *σ*_*L*_, and the covariance between samples, defined by *K*_*γ*_ :

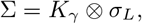

where *⊗* is the Kronecker product: each element of *K*_*γ*_ is multiplied by the matrix *σ*_*L*_ to obtain Σ. The covariance between features from different samples, *K*_*γ*_, has a block-diagonal form, because its elements are zero for features from different participants. For samples from the same participant, the values depend on the time between the samples:

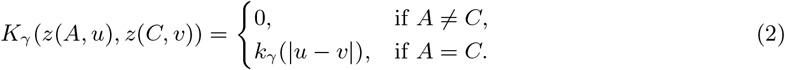

We consider two alternative options for the kernel *k*_*γ*_ : the radial basis function (*RBF*) kernel

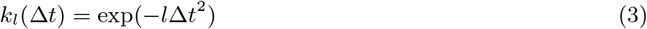

or the *Matérn* kernel

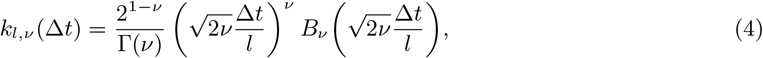

where exp is the exponential function, Γ is the gamma function, and *B*_*ν*_ is the modified Bessel function of the second kind. The kernel parameter *γ* is therefore either *l* or (*l, ν*). The Matérn kernel takes a simpler form for specific values of *ν*; for instance, the Matérn kernel with *ν* = 1*/*2 is known as the exponential kernel:

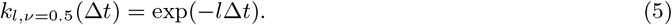

We use a decomposition by a lower-triangular matrix *L* to construct the within-sample covariance in *full-rank* form:

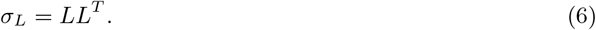

Alternatively, a *low-rank* representation reduces the number of parameters:

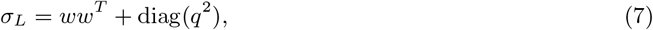

where *w* and *q* are vectors of size *D −* 1 and diag(*q*^2^) is a diagonal matrix with the elements of *q*^2^ on its diagonal.

#### The last value

There are *D −* 1 degrees of freedom in compositional space, because the sum of the features is equal to one. This property allows us to model latent vectors of size *D −* 1 and then append a last value *z*_*D*_ to them. If the appended value is *z*_*D*_ = 0, we call this an *additive* transformation. If it is the negative sum of the vector’s values,

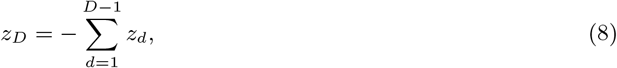

we call it a *centered* transformation; the names follow the additive and centered log-ratios [39]. We denote the resulting vector of size *D* by 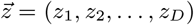.

**Activation functions** transform data from latent space to compositional space. The most common choice is the *softmax* function [36], which exponentiates the elements of 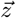 and then normalizes them:

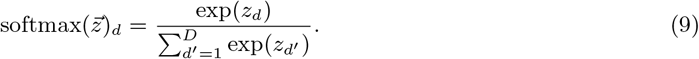

An alternative to softmax, *sparsemax* [40], allows the output compositional vector to contain zero values:

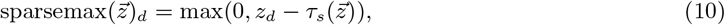

where 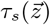 is a threshold determined by the vector 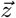 and max(0, *x*) replaces negative *x* by zero.

Both functions belong to a family of activation functions called *a*-entmax [37], which includes softmax when *a* = 1 and sparsemax when *a* = 2. An exact form is also obtained in [37] for *a* = 1.5:

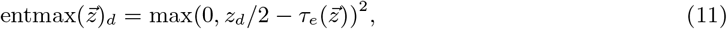

where the threshold 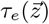 is determined by 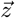. Hereafter, entmax refers to the case *a* = 1.5. The outputs of softmax, sparsemax, and entmax all lie in compositional space.

#### Time offsets

Timestamps may be given as measurement times (e.g., days or months) or as a discrete ascending sequence (e.g., 0, 1, 2, …). However, the distance between timestamps (Δ*t* = |*u − v*| in Eq. 2) may not reflect the difference in features between measurements. For example, a significant event, such as childbirth or an experimental intervention, may occur between timestamps. For this reason, we introduce parameters *t*^*o*^ that shift the original timestamps *t*_1_, *t*_2_, …, *t*_*T*_ . Only *T −* 2 parameters are introduced, as two degrees of freedom correspond to the shift and scale of the timestamps:

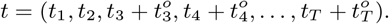

#### Dirichlet distribution

The output *p* of the activation function is taken as the output of the model. Optionally, it can be passed to a Dirichlet distribution [31] together with a scale parameter *α*:

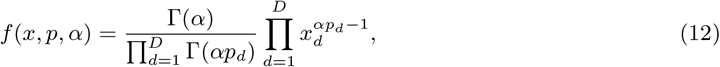

where *f*(*x, p, α*) is the Dirichlet probability density function and *x* lies in compositional space. The scale parameter *α* can also be used as a temperature parameter [41], i.e., as a multiplier of the latent vectors. While this is an explicit way of adjusting the values of the probability vector, the parameter then becomes redundant, as the same scaling can be achieved through *W,b*, and *σ*_*L*_.

### 2.3 Optimization

The optimal parameters of the model are found by a limited-memory algorithm for bound-constrained optimization [42]. To define initial values for the optimization, we transform the compositional vectors in the training set to latent space. This transformation is the inverse of softmax (Eq. 9), sparsemax (Eq. 10), or entmax (Eq. 11). Since all elements are positive, as discussed in Section 2.1, the inverse is the natural logarithm, the identity function, or twice the square root, respectively. We then subtract either the last element or the mean of all elements, to satisfy the additive or centered transformation, respectively, and drop the last dimension. Let *y*^*z*^ denote the latent representation of a vector *y*. The initial values of the parameters, together with their dimensionalities, are given in Table 1.

**Table 1:** Parameters of the model, their dimensionality, and a brief description of their initialization. **Y**^**z**^ is the matrix of stacked vectors *y*^*z*^, and ***β*** is the matrix of all covariates in the training set, stacked in the same order.

| Parameter | dimensionality | initial value |
| --- | --- | --- |
| $b$ | $D - 1$ | mean of $\mathbf{Y}^z$ |
| $W$ | $n_\beta(D - 1)$ | solution of $W\beta = \mathbf{Y}^z - b$ |
| $L$ | $D(D - 1)/2$ if full rank<br>$2(D - 1)$ if low rank | Cholesky decomposition [43] of covariance matrix of $\mathbf{Y}^z - W\beta - b$ if full rank |
| $\gamma$ | 1 or 2 | $l = 1, \nu = 0.5$ |
| $t^o$ | $T - 2$ | 0, 0, ..., 0 |
| $\alpha$ | 1 | 1 |

The optimal parameters of the model are found by minimizing one of four candidate loss functions. All losses are averaged across positions,

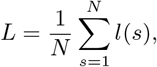

where *l*(*s*) is a function defined for each position in the training set. The first two choices are the negative log-likelihoods of the *Gaussian* distribution (Eq. 1) and the *Dirichlet* distribution (Eq. 12). The third is the mean squared error (*MSE*):

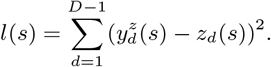

Lastly, we consider the *logistic* loss [37, 40], which has the general form

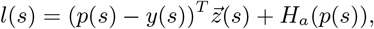

where *H*_*a*_ is the Tsallis entropy [44], a generalization of the Shannon entropy [45], with *a* = 1, *a* = 1.5, and *a* = 2 for softmax, entmax, and sparsemax, respectively.

The Dirichlet and logistic losses use compositional vectors explicitly, while the Gaussian loss and MSE operate on latent representations. The Gaussian loss does not require sampling during training. For all other losses, samples are drawn from the Gaussian distribution with a fixed random seed, and the loss is averaged across all generated samples.

### 2.4 Metrics

We use two metrics to evaluate imputation accuracy, each measuring the distance between a real sample *x* and a predicted sample *y*. The first is the Aitchison distance Δ_*A*_ [39], which is the Euclidean distance in real space after the centered log-ratio transformation (the inverse of softmax satisfying Eq. 8):

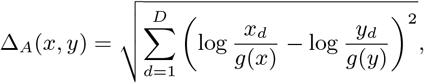

where 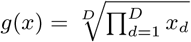 is the geometric mean.

The second is the Jensen–Shannon distance Δ_*JS*_ [46], which is based on a symmetric version of the Kullback–Leibler divergence, KL [47], and measures the dissimilarity between two compositional vectors:

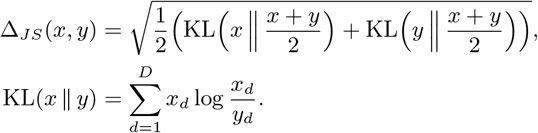

We train the model on the training set and estimate the metrics on the validation set using 5-fold cross-validation. Participants with only one sample are not placed in the validation set for imputation. Samples in the validation set are predicted using the posterior mean and posterior variance of the Gaussian process [28]. For *single* imputation, the prediction is based on the posterior mean. For *multiple* imputation, 100 samples are generated per position. The results are reported with 95% confidence intervals based on Student’s *t*-distribution [48].

### 2.5 Classification

For classification, participants are divided into training and test sets using 5-fold cross-validation. Participants without labels are not involved in training or testing the classifier. In the main text, test sets contain only participants with no missing samples; in this way, we isolate testing from the effect of imputation.

Classification performance is measured by balanced accuracy, the mean of sensitivity and specificity. Sensitivity is the true positive rate, i.e., the fraction of participants with a positive label that are correctly classified. Specificity is the true negative rate, i.e., the fraction of participants with a negative label that are correctly classified.

For each of 10 imputed datasets, we perform the following steps for binary classification:

0. Optional log-ratio transformation: we take the ratios of features at each time point and use the logarithms of these ratios.
1. Feature selection: a subset of features is selected based on their positive coefficients in an elastic net [49]. Elastic-net regularization penalizes the coefficients of a logistic regression that classifies the labels, thus highlighting more meaningful and independent features. The features are selected on validation sets (part of the training set, not the test set).
2. Classification: we apply an extremely randomized trees classifier [50] to predict labels from the constructed feature space.

## 3 Datasets

We train and test the model on three datasets.

**The BASIC study** [51] collected gut microbiome samples from 512 pregnant women at Uppsala University Hospital from 2016 to 2019. Samples were collected at *T* = 3 time points: during the 20th and 30th weeks of pregnancy and at six weeks postpartum. Samples with fewer than 500,000 shotgun metagenomic reads were excluded from the analysis. Raw reads were quality-controlled and annotated with the MetaPhlAn pipeline as previously described [52]. In total, 937 microbiome samples were available for analysis, corresponding to 61% of all possible samples.

Depressive symptoms were assessed at each visit using the Edinburgh Postnatal Depression Scale (EPDS) questionnaire [53]. The prediction target is depression status at the postpartum visit, where participants were classified as healthy if their EPDS score was below 12 and as depressed otherwise [54]. The covariate vector *β* has size *n*_*β*_ = 7: it includes the timestamp, a 3-dimensional vector indicating whether the EPDS score at each time point is *{*below 12, 12 or above, or missing*}*, and a 3-dimensional vector, shared across all time points for a given participant, indicating which sampling visits were completed.

We selected the two bacterial species whose log-ratio maximized the ANOVA *F*-statistic [55] for separating healthy and depressed participants [56]: (1) *Haemophilus parainfluenzae* and (2) *Veillonella parvula*, with component *d* = *D* = 3 aggregating the remaining species. To extend the analysis to higher dimensionalities (*D* = 4 and *D* = 9), additional subsets were chosen via permutation importance evaluated across a candidate pool of 12 depression-associated taxa [57–61]: (3) *Bilophila wadsworthia*, (4) *Ruminococcaceae* bacterium D16, (5) Clostridiales bacterium 1747FAA, (6) *Akkermansia muciniphila*, (7) *Prevotella histicola*, (8) Proteobacteria bacterium CAG 139, (9) Clostridiales bacterium CHKCI006, (10) *Ruminococcaceae* bacterium D5, (11) *Faecalibacterium prausnitzii*, (12) *Parasutterella excrementihominis*. The subsets (3–5) and (2–9) comprise the taxa with positive permutation importance at all time points and at one or more time points, respectively. The importance is obtained by randomly permuting the values of a feature and quantifying the resulting decrease in balanced accuracy.

### The GMAP study

As part of the Gastrointestinal Microbiome and Allergic Proctocolitis (GMAP) study, longitudinal samples from infants up to the age of one year with food protein-induced allergic proctocolitis (FPIAP) and from matched controls were analyzed [62]. We consider three time points in the first half of the first year, namely months 2, 4, and 6. There are 157 infants with at least one sample at these three time points. The percentages of missing samples per time point are 21%, 33%, and 38%, respectively. The results are shown for the four bacterial genera with the largest ANOVA *F*-statistic [56] for distinguishing between cases and controls: *Lactobacillus, Dorea, Blautia*, and *Veillonella*.

There are *n*_*β*_ = 8 covariates for this dataset: three indicate whether an appointment was missed, and the fourth is the month of the visit. The other four are binary variables representing the child’s sex, the mode of delivery, whether the infant was the first child in the family, and whether breastfeeding was initiated.

### The Wellness study

The third dataset is taken from the Swedish SciLifeLab SCAPIS Wellness Profiling study [63]. We consider gut microbiome data from *T* = 4 visits of *n* = 101 individuals aged 50–65 years, with 384 available samples in total. We select the seven species with the lowest and the seven with the highest total variance according to [64], thus constructing two datasets with *D* = 8: (*Clostridium symbiosum, Holdemania filiformis, Subdoligranulum variabile, Ruminococcus obeum, Coprococcus comes, Faecalibacterium prausnitzii, Roseburia inulinivorans*, and the abundance of the remaining species) and (*Methanobrevibacter smithii, Dialister invisus, Desulfovibrio piger, Akkermansia muciniphila, Sutterella wadsworthensis, Prevotella copri, Escherichia coli*, and the abundance of the remaining species).

While the dimensionality of the feature space is not a limitation of the model itself, we keep the number of parameters below the number of available samples so as not to sacrifice statistical accuracy or computational efficiency [65]. The covariates are constructed by concatenating the timestamp with four binary variables denoting missing samples; thus *n*_*β*_ = 5.

## 4 Results

### Study 1

We evaluated the GRF model configurations against the LOCF baseline [35] using the *D* = 3 feature subset of the BASIC dataset. The GRF was tested across activation and loss functions (Table 2). The two metrics (Aitchison and Jensen–Shannon) were consistent with each other: lower values of Δ_*A*_ corresponded to lower values of Δ_*JS*_ . The model using entmax trained with the logistic loss and the model using softmax trained with the Gaussian or MSE loss outperformed LOCF. The GRF with entmax activation produced samples closest to the real samples in the validation sets. The GRF is also able to perform multiple imputation, reflecting the uncertainty around the imputed values [16]. Softmax with the MSE loss and entmax with the logistic loss achieved a low error for multiple imputation.

**Table 2:** Comparison of imputation methods for the BASIC study with *D* = 3. Last (next) observation carried forward (backward) is compared to the GRF with an exponential kernel, a full-rank within-sample covariance, and an additive transformation. For the GRF, three activation functions and four loss functions are presented. Results are given for single and multiple imputation under two metrics: the Aitchison distance and the Jensen–Shannon distance (multiplied by 10). Each distance is given with its 95% confidence interval. * denotes cases where zero values produced by the model are replaced by the value *ϵ*. Bold values are the minimum in each column.

| activation | loss | Aitchison ($\Delta_A$) | | Jensen–Shannon ($\Delta_{JS}$) $\times 10$ | |
| --- | --- | --- | --- | --- | --- |
|  |  | single | multiple | single | multiple |
| Last/Next carried |  | 2.76(2.6 2.92) | - | 0.09(0.09 0.1) | - |
| softmax | Gaussian | **2.69**(2.6 2.78) | 3.68(3.46 3.9) | **0.08**(0.07 0.09) | 0.15(0.11 0.19) |
|  | Dirichlet* | 5.33(5.04 5.62) | 4.41(4.14 4.69) | 0.72(0.68 0.76) | 0.35(0.24 0.45) |
|  | logistic | 3.03(2.82 3.23) | 3.03(2.88 3.17) | 0.09(0.09 0.1) | 0.09(0.09 0.1) |
|  | MSE | 2.73(2.65 2.81) | 2.73(2.67 2.78) | **0.08**(0.07 0.09) | **0.08**(0.08 0.09) |
| sparsemax* | Gaussian | 3.52(3.27 3.76) | 4.5(4.23 4.78) | 0.12(0.12 0.12) | 0.22(0.2 0.25) |
|  | logistic | 3.59(3.3 3.87) | 3.59(3.4 3.79) | 0.12(0.11 0.12) | 0.12(0.12 0.12) |
|  | MSE | 3.53(3.26 3.8) | 3.53(3.35 3.71) | 0.12(0.12 0.12) | 0.12(0.12 0.12) |
| entmax* | Gaussian | 2.98(2.85 3.11) | 3.82(3.59 4.05) | 0.09(0.08 0.09) | 0.13(0.11 0.14) |
|  | logistic | **2.7**(2.52 2.87) | **2.7**(2.57 2.82) | **0.08**(0.07 0.09) | **0.08**(0.07 0.09) |
|  | MSE | 3.02(2.87 3.16) | 3.02(2.92 3.12) | 0.09(0.08 0.09) | 0.09(0.08 0.09) |

Real samples and samples from single imputation across all validation sets are plotted in Figure 3 for each activation function. The figure shows that entmax and sparsemax reflected the sparsity of the data better than softmax, while softmax and entmax yielded a lower distance between predictions and target samples.

**Figure 3:**
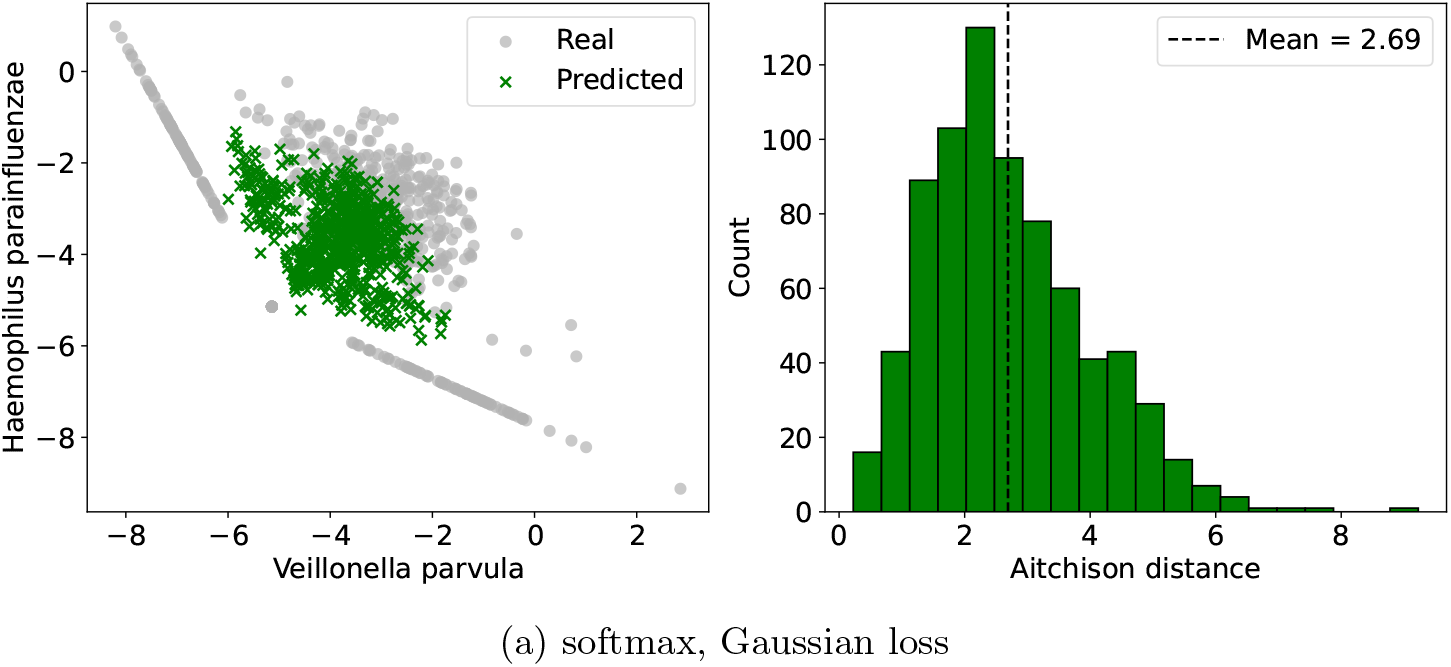
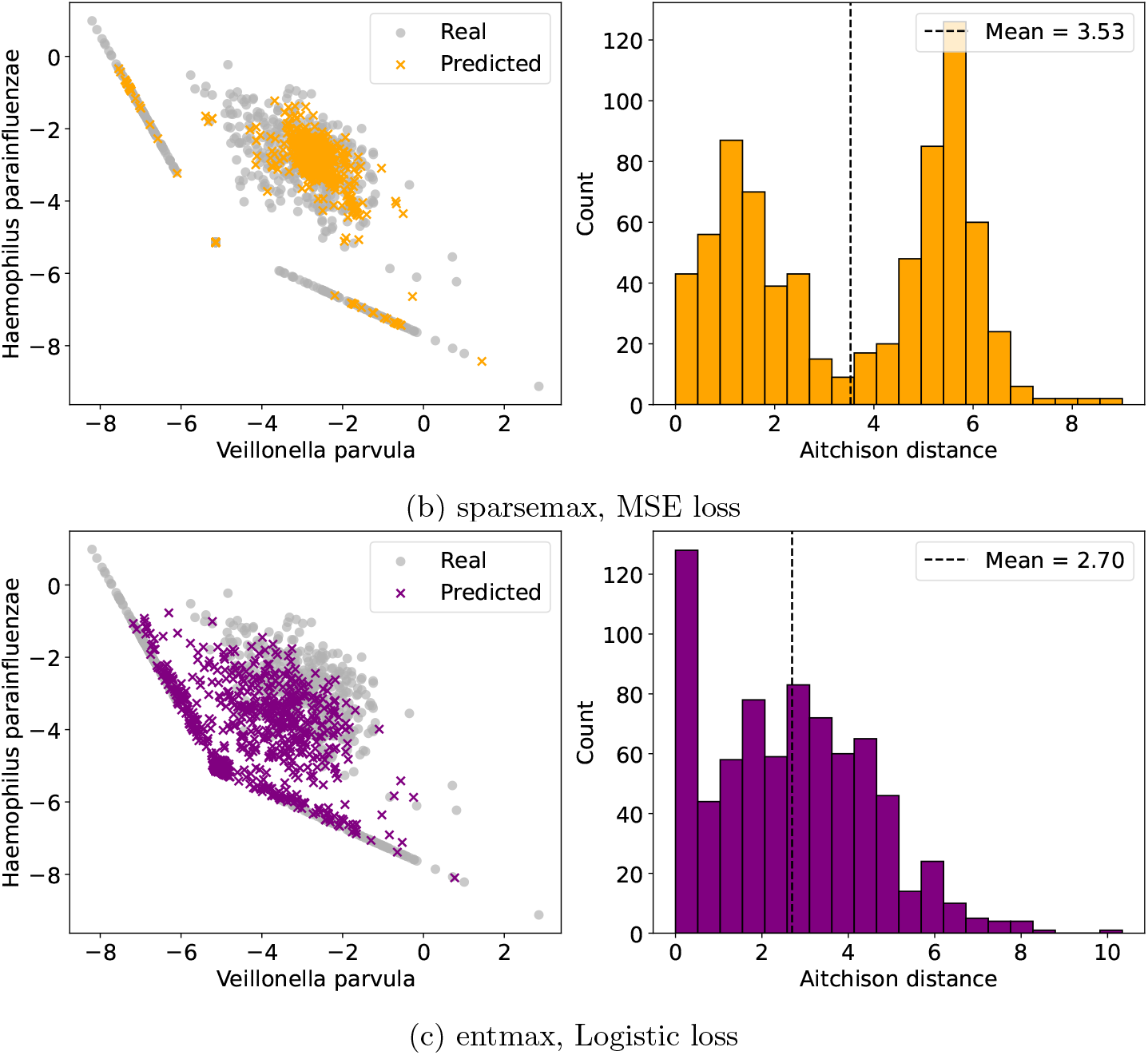
Left: real samples from the BASIC study (*D* = 3) and samples predicted by the trained GRF model, shown across all cross-validation folds. Samples are presented in two dimensions after the centered log-ratio transformation. The two dotted (gray) lines correspond to samples in which a bacterial feature has the minimum abundance. Right: histograms of Aitchison distances between real and generated samples. (a) Model with softmax activation trained with the Gaussian loss. (b) Model with sparsemax activation trained with the MSE loss. (c) Model with entmax activation trained with the logistic loss.

The primary model configuration used an exponential kernel, a full-rank within-sample covariance matrix, and an additive transformation (*z*_*D*_ = 0). To complete the analysis of imputation in the three-feature space, Table 3 presents alternative configurations of the model. The GRF model is sensitive to the choice of kernel in the Gaussian process. A statistically insignificant decrease in imputation error was obtained with the two-parameter Matérn kernel. The choice of covariance matrix and of the transformation in latent space had no significant effect.

**Table 3:**
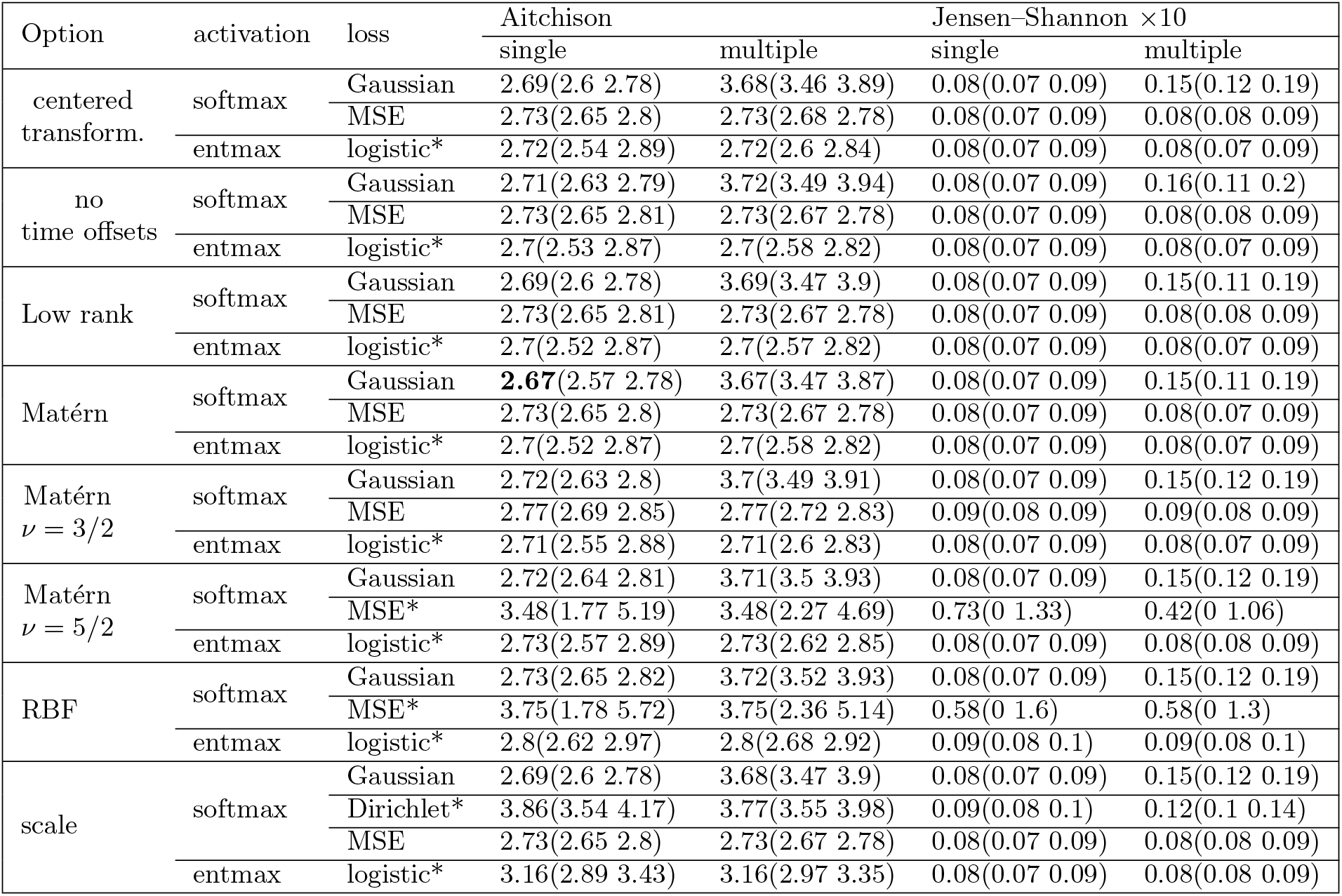
Comparison of the GRF model variations described in Section 2.2 for the BASIC study with *D* = 3. The activation and loss functions are given in the second and third columns. Results are given for single and multiple imputation under two metrics: the Aitchison distance and the Jensen–Shannon distance (multiplied by 10). Each distance is given with its 95% confidence interval. The options, from the top, are: the centered transformation for the last value in latent space (Eq. 8), time offsets set to zero, the low-rank within-sample covariance (Eq. 7), the Matérn kernel (Eq. 4) with *ν* as a parameter of the model and with *ν* = 3*/*2 or *ν* = 5*/*2, the RBF kernel (Eq. 3), and inclusion of the scale parameter *α*. * denotes cases where zero values produced by the model are replaced by the value *ϵ*. Values are in bold if they are lower than those obtained with the default options: exponential kernel, full-rank within-sample covariance, and additive transformation.

We selected the activation and loss functions based on imputation accuracy and tested them in a downstream analysis. We used multiple imputation to reconstruct all missing values and used the imputed datasets for a forecasting task. After imputation, we considered the two time points during pregnancy and aimed to forecast depression at the last time point, *t* = 3, postpartum. We constructed the input space by concatenating the microbiome features with two covariates on the depression state at *t* = 2, denoting the EPDS value and its presence or absence in the collected data. The results are shown in Table 4, which compares compositional inputs from the microbiome with their log-ratios. Completing the dataset with the GRF model with entmax activation, followed by the classification pipeline, yielded a balanced accuracy of 0.67. Repeating the imputation and forecasting procedure with different random seeds, we obtained an estimated balanced accuracy of 0.64, a slight improvement over the previous result of 0.62 based on the same bacteria [56]. However, forecasting ability did not increase when additional species were included in the input space.

**Table 4:**
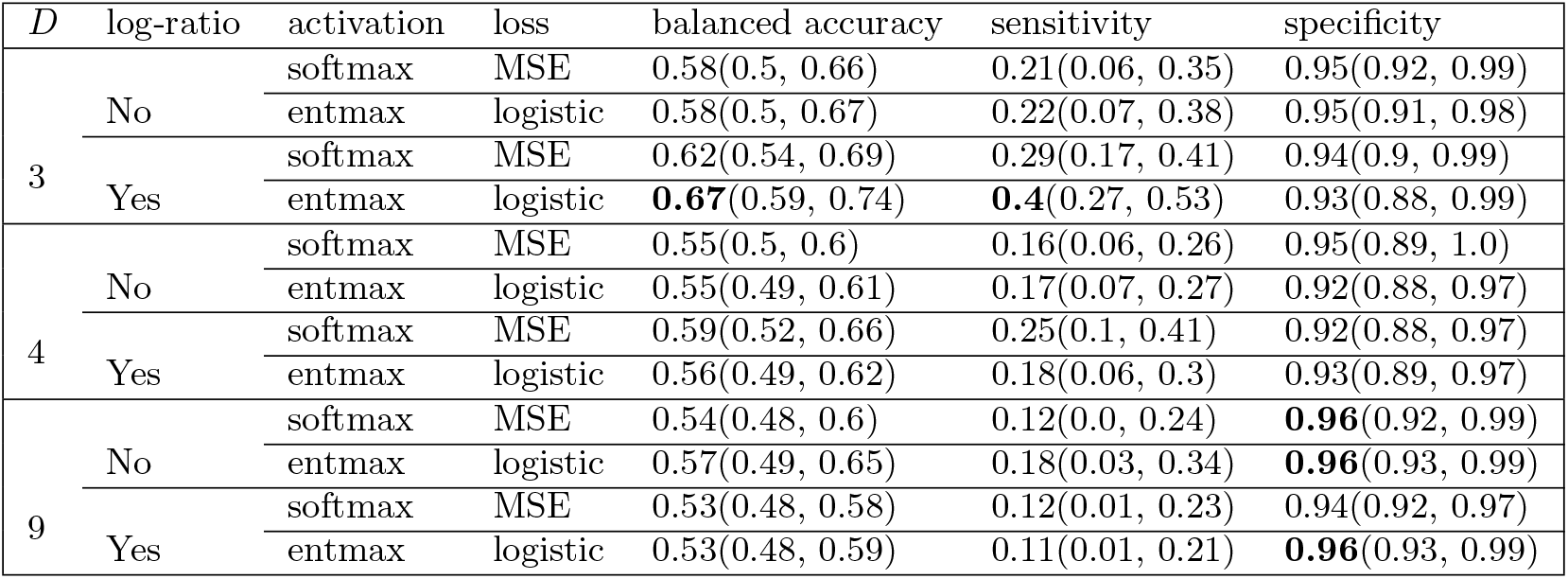
Balanced accuracy, sensitivity, and specificity for forecasting postpartum depression from the gut microbiome with *D* = 3, 4, and 9 relative abundances. A low-rank approximation of the covariance matrix was used for *D* = 9. The second column indicates whether the log-ratio transformation was applied to the compositional samples before forecasting. Multiple imputation was performed using the GRF model with the activation function in column 3, trained with the loss function in column 4. Bold values are the maximum in each column.

We also ran experiments with missing data in the test set, obtaining a balanced accuracy of 0.58 with a sensitivity of 0.66. This corresponds to correctly identifying two thirds of the depressed women, but having only a 50% chance of correctly classifying a healthy participant. These and additional results for *D* = 3 are given in Appendix A.

### Study 2

As an additional test, we performed binary classification of cases and controls in the GMAP study of food protein-induced allergic proctocolitis (Table 5). Whereas we previously reached a balanced accuracy of 0.66 when using all study participants [56], here we achieved a balanced accuracy of 0.72 with the entmax function. Although the two loss functions yielded the same balanced accuracy, the MSE loss was balanced in sensitivity and specificity, whereas the logistic loss gave the best sensitivity at the expense of lower specificity. The balanced accuracy obtained with the different activation functions corresponds to the imputation accuracy of the respective GRF configurations (Figure 4). The Aitchison distance obtained with LOCF is 6.97 in this case.

**Table 5:** Balanced accuracy, sensitivity, and specificity for forecasting FPIAP from the gut microbiome in the GMAP study. A low-rank approximation of the covariance matrix was used. Features were extracted from the compositional samples before forecasting. Multiple imputation was performed using the GRF model with the activation function in column 1, trained with the loss function in column 2. Bold values are the maximum in each column.

| activation | loss | balanced accuracy | sensitivity | specificity |
| --- | --- | --- | --- | --- |
| softmax | MSE | 0.58(0.43, 0.73) | 0.51(0.23, 0.79) | 0.66(0.46, 0.85) |
|  | logistic | 0.64(0.54, 0.74) | 0.62(0.44, 0.79) | 0.66(0.52, 0.8) |
| entmax | MSE | **0.72**(0.61, 0.83) | 0.74(0.63, 0.84) | **0.71**(0.51, 0.91) |
|  | logistic | **0.72**(0.58, 0.87) | **0.8**(0.66, 0.94) | 0.64(0.4, 0.89) |

**Figure 4:**
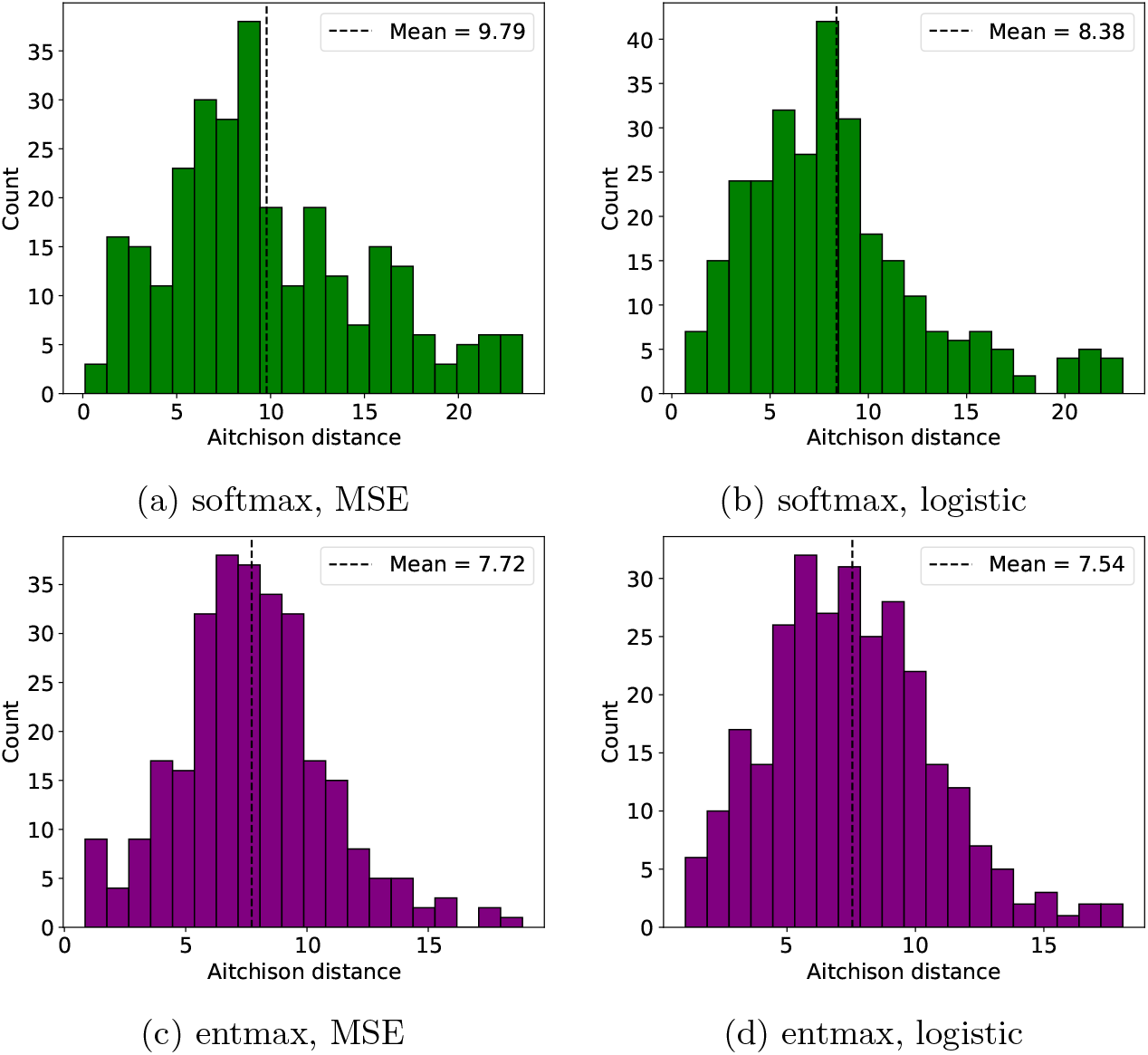
Histograms of Aitchison distances between real and generated samples in the GMAP study, for the model with softmax activation (top row) or entmax activation (bottom row), trained with the MSE loss (left column) or the logistic loss (right column).

### Study 3

The Wellness study does not include an outcome-related set of bacteria that could be analyzed. We therefore tested the limits of the model by applying it to either the least or the most variable species in the dataset. Applying the GRF model to the least variable species, we obtained a low Aitchison distance with the softmax activation and a low Jensen–Shannon distance with the entmax activation trained with the MSE loss (Table 6). In contrast to the first dataset, sparsemax gave a low Jensen– Shannon distance for all loss functions. However, for the species with the highest variance (Table 7), we were unable to achieve the same accuracy as sample carrying, likely owing to less informative covariates and a smaller training dataset than in the other studies.

**Table 6:** Comparison of imputation methods for the species with low total variance in the Wellness study. Last (next) observation carried forward (backward) is compared to the GRF with an exponential kernel, a full-rank within-sample covariance, and an additive transformation. For the GRF, three activation functions and four loss functions are presented. Results are given for single and multiple imputation under two metrics: the Aitchison distance and the Jensen–Shannon distance (multiplied by 10). Each distance is given with its 95% confidence interval. * denotes cases where zero values produced by the model are replaced by the value *ϵ*. Bold values are the minimum in each column.

| activation | loss | Aitchison | | Jensen–Shannon $\times 10$ | |
| --- | --- | --- | --- | --- | --- |
|  |  | single | multiple | single | multiple |
| Last/Next carried |  | **3.58**(3.26 3.9) | - | 0.64(0.6 0.68) | - |
| softmax | Gaussian | 5.27(4.26 6.27) | 6.74(6.11 7.38) | 1.06(0.66 1.46) | 1.69(1.46 1.93) |
|  | Dirichlet* | 12.56(11.95 13.18) | 12.16(11.75 12.57) | 2.57(2.53 2.6) | 3.51(3.21 3.81) |
|  | logistic | 4.72(4.37 5.07) | 4.88(4.68 5.07) | 0.64(0.56 0.72) | 0.64(0.58 0.7) |
|  | MSE | 4.02(3.81 4.23) | **4.02**(3.87 4.17) | 0.65(0.6 0.7) | 0.65(0.62 0.69) |
| sparsemax* | Gaussian | 6.12(5.95 6.28) | 9.67(9.17 10.18) | 0.63(0.59 0.66) | 0.92(0.86 0.97) |
|  | logistic | 5.63(4.5 6.77) | 6.14(5.64 6.63) | 0.63(0.59 0.66) | 0.63(0.61 0.66) |
|  | MSE | 6.24(5.99 6.49) | 6.26(5.93 6.59) | 0.63(0.59 0.66) | 0.63(0.6 0.65) |
| entmax* | Gaussian | 6.14(5.97 6.31) | 6.98(6.58 7.38) | 0.73(0.66 0.79) | 0.9(0.85 0.96) |
|  | logistic | 5.82(5.4 6.25) | 5.89(5.62 6.17) | 1.24(1.18 1.29) | 1.24(1.2 1.28) |
|  | MSE | 6.18(5.98 6.38) | 6.18(6.03 6.32) | **0.61**(0.58 0.65) | **0.61**(0.59 0.64) |

**Table 7:** Comparison of imputation methods for the species with high total variance in the Wellness study. Last (next) observation carried forward (backward) is compared to the GRF with an exponential kernel, a full-rank within-sample covariance, and an additive transformation. For the GRF, three activation functions and four loss functions are presented. Results are given for single and multiple imputation under two metrics: the Aitchison distance and the Jensen–Shannon distance (multiplied by 10). Each distance is given with its 95% confidence interval. * denotes cases where zero values produced by the model are replaced by the value *ϵ*. Bold values are the minimum in each column.

| activation | loss | Aitchison | | Jensen–Shannon $\times 10$ | |
| --- | --- | --- | --- | --- | --- |
|  |  | single | multiple | single | multiple |
| Last/Next carried |  | **5.63**(5.29 5.96) | - | **0.86**(0.78 0.94) | - |
| softmax | Gaussian | 17.45(14.31 20.59) | 19.44(17.31 21.56) | 3.71(3.07 4.34) | 4.18(3.72 4.65) |
|  | Dirichlet* | 10.86(9.72 12.0) | 12.56(11.95 13.18) | 2.21(1.84 2.57) | 2.78(2.44 3.12) |
|  | logistic | 9.95(8.66 11.25) | 9.95(9.04 10.87) | 1.12(0.93 1.31) | 1.12(0.98 1.25) |
|  | MSE | 8.42(8.08 8.76) | 8.42(8.18 8.66) | 1.37(1.28 1.47) | 1.37(1.31 1.44) |
| sparsemax* | Gaussian | 14.28(13.64 14.93) | 15.7(15.07 16.33) | 1.41(1.33 1.49) | 1.63(1.56 1.71) |
|  | logistic | 12.88(12.75 13.02) | 12.89(12.78 12.99) | 1.3(1.25 1.35) | 1.3(1.26 1.33) |
|  | MSE | 12.75(12.65 12.86) | 12.8(12.67 12.94) | 1.3(1.25 1.35) | 1.3(1.27 1.33) |
| entmax* | Gaussian | 13.51(12.87 14.14) | 14.0(13.41 14.58) | 1.4(1.27 1.53) | 1.58(1.49 1.67) |
|  | logistic | 10.06(9.73 10.39) | 10.06(9.83 10.29) | 1.14(1.1 1.19) | 1.14(1.11 1.17) |
|  | MSE | 11.83(11.67 12.0) | 11.83(11.71 11.95) | 1.1(1.05 1.16) | 1.1(1.06 1.14) |

## 5 Discussion

We have presented an explainable GRF framework for the multiple imputation of longitudinal compositional data. The model allows us to reconstruct time series of features and to increase the number of complete data points. It addresses all three dimensions of longitudinal studies: a set of participants with different characteristics and collected data, measurements taken repeatedly over time for each participant, and internal relationships among the measurements. The model is flexible with respect to the number of participants, time points, and features, and we have applied it to data varying in the number of features and in the extent of temporal variation. Its moderate number of interpretable parameters provides a computationally efficient alternative to existing generative neural architectures [20, 25, 26], while decreasing the risk of overfitting.

Generating realistic samples in a longitudinal setting, with several compositions available per participant, is particularly challenging because of the dual constraints of a multi-dimensional compositional feature space and participant-dependent temporal dynamics. The core strength of our approach lies in overcoming this challenge and generating time-dependent synthetic compositions. By explicitly modeling the joint covariance across latent features, our model generates unobserved states that capture dynamic biological shifts and multivariate co-dependencies. To transform real vectors into compositional space, we couple the GRF with activation functions, which allows the model to produce zero values naturally and to reflect data sparsity. Although copying the nearest observed sample guarantees biologically real data, our approach achieves comparable or superior imputation accuracy.

We implemented the GRF model with a flexible choice of architecture. We performed a comparative analysis of activation functions and proposed a variety of loss functions for training the model. We observed that the choice of kernel influences imputation accuracy. We also extended the GRF model by introducing parameters corresponding to shifts of the timestamps (Fig. 2). The design of the model allows us to differentiate between distinct types of missingness in the data; for instance, we mark missed appointments in a participant’s history, but not samples missing because of insufficient reads. The choice of activation function mapping to compositional space regulates sample sparsity. Although the optimal model configuration remains data-dependent, we recommend pairing the GRF with the 1.5-entmax activation as the default option, as it yielded superior average imputation performance across the evaluated datasets. An implementation of the model, for reproducibility and further use, is available in a GitHub repository^1^.

The probabilistic formulation provides multiple imputations, allowing incomplete longitudinal data to be resolved into an ensemble of completed datasets suitable for downstream analyses such as classification or forecasting. To demonstrate this, we investigated the forecasting ability of imputed gut microbiome samples in two datasets with participant-level labels. Applying a classification pipeline to the imputed datasets under repeated cross-validation, we reached a balanced accuracy of 0.72 for classifying infants with food protein-induced allergy at 12 months. This result was obtained by reconstructing missing samples for infants at three time points (months 2, 4, and 6). The proposed framework with multiple imputation by the GRF improved both sensitivity and specificity compared with the original study of these data [62], which used additional data points, and compared with our previous study of the same data, which achieved a balanced accuracy of only 0.66 [56]. Likewise, applying a similar pipeline to samples collected during pregnancy to forecast postpartum depression, we achieved a balanced accuracy of 0.64. Table 8 summarizes the improvements in balanced accuracy and sensitivity relative to previous studies [56] when assigning all incomplete data to the training set prior to imputation.

**Table 8:** Downstream classification performance when retaining and imputing incomplete trajectories in the training set, benchmarked against [56] (left values in columns). Bold values indicate the improved results achieved in this work.

| Target Outcome | Balanced Accuracy | Sensitivity |
| --- | --- | --- |
| Postpartum depression | 0.61 → **0.64** | 0.25 → **0.34** |
| Food allergy | 0.66 → **0.72** | 0.65 → **0.80** |

The model assumes that all participants are independent, which allows for a faster implementation, since the covariance matrix has a block-diagonal form. This assumption can be relaxed, at the cost of training speed, by incorporating inter-participant distances, e.g., if participants in a study come from different geographical locations or are relatives. The model relies on the covariates known about the participants and on the timestamps. A limitation of this approach is its dependence on how informative these variables are for constructing the imputations, which is why the model does not perform as well on the Wellness data as on the other datasets.

The model uses information gained from the available samples to impute compositional vectors in place of missing samples. Its application can be extended to data augmentation, i.e., generating samples at all time points for artificial participants. Data augmentation helps to balance data for downstream analysis; for example, women with postpartum depression represent a minority in the BASIC study. The model can also be applied to the test set if it contains missing samples: in this scenario, we attained a balanced accuracy of 0.58 with a high sensitivity of 0.66.

The model is specifically constructed for datasets with a small to moderate number of samples and therefore includes a limited number of parameters. It is, however, scalable to larger datasets: estimating the mean in latent space and applying the activation function can be viewed as a single layer of a neural network. Future extensions of the model may place prior distributions on the model parameters, as done in [33], while the parameter *a* of *a*-entmax [37] could be learned directly by the model instead of being chosen from the discrete set *{*1, 1.5, 2*}*. In summary, we have presented a scalable, efficient, and accurate model for imputing missing data in compositional time series, which can improve the power and analytic flexibility of any kind of biological time-series analysis in the future.

### A Forecasting

We conducted additional experiments with *D* = 3, the log-ratio transformation, entmax activation, and the logistic loss.

- The EPDS covariates at time point 1 alone are not informative: the balanced accuracy with only these two features is 0.5.
- Providing information about missingness to the classifier does not improve performance: the balanced accuracy is 0.66 (0.57, 0.75) when two covariates about missing samples are added to the input space.
- Including participants with missing samples in the test set decreases the balanced accuracy to 0.57 (0.52, 0.61). Missingness in the test set requires changes to the training of the imputation model: all samples at *t* = 3 were excluded, and information about missing samples at *t* = 3 was removed from the covariates.
- We repeated the experiment from the previous item with a different data balance: in the training set, time points were imputed only for participants with a positive label. The balanced accuracy then becomes 0.58 (0.52, 0.63), while the sensitivity increases to 0.66 (0.59, 0.73), with a specificity of 0.49 (0.43, 0.55). We applied the same balancing approach in [56], whereas in Section 4 we use all available participants in the training set.
- We extended the experiment in Table 4 to five random seeds for cross-validation, the GRF, and the classifier, attaining a balanced accuracy of 0.64 (0.62, 0.67) with a sensitivity of 0.34 (0.3, 0.39).

## Footnotes

1 https://github.com/Shternshis/GRF_impute

